# Whole-body positive synchronization pattern of metabolic transcriptomes

**DOI:** 10.1101/2023.11.16.567365

**Authors:** Ehud Z. Sussman, Dana Mor, Judith Somekh

## Abstract

Understanding the relationship between transcriptomes in the entire body is crucial for comprehending health and disease. To achieve this, we have developed a framework that allows us to measure the general coordination pattern of functionally similar transcriptomes at a whole-body scale. Our approach employs meta-network level analysis to determine the interconnectivity level of metabolic co-expression modules throughout the body as opposed to randomly selected modules. We utilize dimensionality reduction of co-expression modules, inter-tissue connectivity, and clique community analysis. We applied our methodology to 19 distinct human tissues obtained from the GTEx project and detected 40/52 metabolic co-expression modules out of 609/652 cross-tissue generated modules in Cohort 1 and 2 respectively. We detected that the ratio of positive to negative inter-tissue co-expression of metabolic modules was significantly higher than control. We further validated these results for an additional cohort. Moreover, to detect a global concurrent synchronization pattern we performed graph theoretical analysis, specifically a clique community analysis of metabolic transcriptomes showing a significantly larger inter-tissue metabolic transcriptome community that control with a significantly higher connectivity for each metabolic module within the community. Additionally, we performed gene-level validation of our results. In summary, our findings reveal a significant global synchronization pattern of metabolic modules throughout the body. Our framework can be further used for other type of transcriptomes and serve as a measure detecting change in this global coordination pattern across conditions.

## 1. Introduction

Whole body synchronization and coordination are essential for maintaining homeostasis, responding to environmental changes, and enabling complex biological functions. They exemplify the intricate nature of biological systems, where various elements must work in harmony to ensure an organism’s survival and well-being. Whole-body physiological homeostasis is pivotal for maintaining human health and longevity and includes concurrent cross-tissue regulation. In parallel to preserving this whole-body homeostasis, these co-regulation mechanisms are required to respond to various signals, including internal and external stimuli, such as glucose intake and circadian rhythms [1]. These internal or external stimuli, such as metabolic signals [2]–[4], as well as circadian clocks [5]–[7] were shown to regulate gene expression. A pivotal example is while in the fight or flight phenomenon, the individual is required to recruit its whole system for immediate running and count on these internal synchronization and coordination mechanisms. This synchronization in the context of running refers to the harmonious coordination of various physiological and biomechanical processes involved in the act of running. These processes work together to optimize efficiency, conserve energy, and achieve effective locomotion. This critical whole-body responsiveness requirement to external events requires dedicated coordination patterns. Although this is an important mechanism, this topic of whole-body coordination patterns was poorly investigated. Currently, with the rise in multi-tissue transcriptomic databases, such as the GTEx, that collected samples of multiple tissues from each individual and include RNA-seq data extracted from various tissues from ∼1000 individuals we can investigate transcriptomic tissues synchronicity at a whole-body level rather than test a single tissue each time. A question arises: How the human whole-body transcriptomes are co-regulated and coordinated? And as a first step to tackle this challenge we ask - can we detect a general synchronization pattern at the whole-body level?

While a single tissue co-expression of genes was extensively studied, the human inter-tissue research is very limited [8]–[10] To test inter-tissue gene connectivity, the research in Dobrin et al. [8] integrated genes from 3 tissues, the hypothalamus, liver, and adipose, of healthy and obese mice and constructed co-expressed modules and showed connectivity between interesting genes and coexpressed modules. Seldin et al. [9] calculated endocrine associations in 6-tissue mouse data between ligands expression levels in source tissues and affected genes that form a pathway in the target tissue, connections that were then validated experimentally. The work of Long et [10] calculated pairwise inter-tissue gene-to-pathway connections in humans using the e Genotype-Tissue Expression project, (GTEx) [11], [12].These studies include pairwise inter-tissue interactions between several tissues and are based solely on direct gene-level correlations between the expression of genes and genes within pathways. Single-cell sequencing technologies add the dimension of heterogeneity beyond tissue type within a single tissue. Furthermore, Mair et al. [13] demonstrated by focusing on a subset of genes associated with the immune system gene expression levels are susceptible to external activation and can vary dramatically over a span of hours. Although these are robust efforts, systematic and system-level studies of tissue interactions are poorly addressed and miss a whole-body system-level perspective. Moreover, these efforts that count on a single gene association with other genes or pathways may be prone to noise affecting the levels of single genes. Indeed, the GTEx data is highly heterogeneous and influenced by external artifacts that need to be corrected [14].

Here, we developed a computational framework to evaluate a global cross-tissue transcriptomic synchronization pattern, which we applied to metabolic transcriptome across 19 human tissues of 2 cohorts. For this challenge, we developed a large-scale inter-tissue system-level meta-network computational framework encompassing community analysis of dimensionality reduction of coexpression modules across tissues. We used co-expression modules and further validated the results at the single gene levels to show a similar trend. Our framework (see Fig. 1) includes the following steps: (1) Co-expression module generation using the Weighted Gene Coexpression Network Analysis (WGCNA) [15] algorithm, (2) Module annotation into eight categories (e.g., Metabolism) using enrichment analysis and plurality votes, (3) Community analysis using the CPM (K-Clique Percolation Method) algorithm2 [16] .We used the Monte-Carlo randomization tests [17] to evaluate the significance level of the degree of co-expression of eigengenes of functionally similar co-expression modules (e.g., metabolic), community size and connectivity across the whole human body. Our community analysis tests the metabolic positive labeled assortativity characteristic [18], [19] of our whole-body network, i.e., that tests if similar modules (metabolic modules) exhibit more robust positive connectivity and larger communities than randomly selected actual modules. Labeled assortativity [18], [19] is the tendency in networks that nodes with the same label (“metabolic”) will have greater interconnectivity. We applied our approach to the metabolic coordination of 19 tissues validated on two human cohorts derived from the GTEx dataset. We further validated the coordination pattern at the gene level rather the module level, using 12 representative tissue-tissue pairs and 1250 metabolic genes.

**Figure 1.**
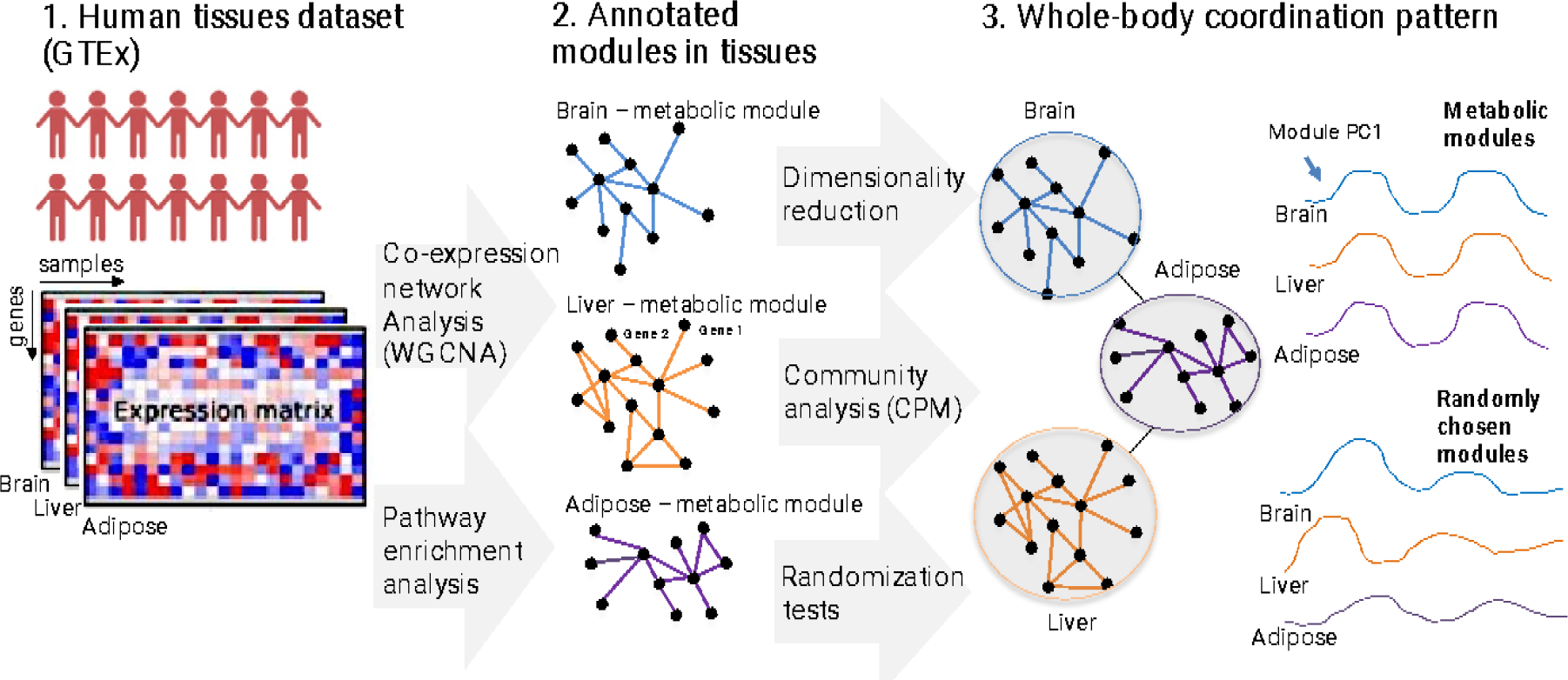
Schematic view and steps of of our methodology.

## 2. Materials and Methods

### 2.1. Data

We downloaded the RNA-seq data from the GTEX project [12]. The GTEx cohort includes postmortem samples extracted from relatively healthy donors at time of death. We limited our study to 19,200 protein-coding genes based on HGNC [20].

### 2.2. Pre-processing

The death classification of the samples (DTHHRDY) was along the Hardy Scale [21] We filtered the samples into two cohorts: (1) Cohort 1 - samples of death types 1 and 2 (violent and fast death or fast death due to natural causes), (2) Cohort 2 - donors that were attached to a ventilation machine before death. We split into two cohorts since we observed from our prior work [22] that mechanical ventilation impacts multiple tissues and is related to ischemic time (the time between death and sample extraction).

We selected the top 19 tissue types (paired tissues sample size > 98 for the smaller cohort 1). We excluded the brain since we wanted to keep a comparable set for both cohorts, and there is a minimal number of brain tissue samples in the second cohort. We log scaled the gene expression levels measured in Transcripts per million (TPM) values, removed ∼2% of outlier samples per tissue, and imputed missing values to nominal values for sample attributes by taking the average of missing SMTSISCH (i.e., ischemic time) for all replicates. For each tissue with a missing value entering the average SMTSISCH value for that particular tissue, longer ischemic times correlate with low-quality samples [23]. We removed tissue-specific genes with minimal sample values, quantile normalized the samples, and corrected for known confounding factors using a multiple linear regression model. We corrected for batch effects, ischemic time (the time that elapsed from the actual time of death to the time of sample extraction), sample quality (most samples include high-quality values of SMRIN > ∼6), gender, and age using the following equation:

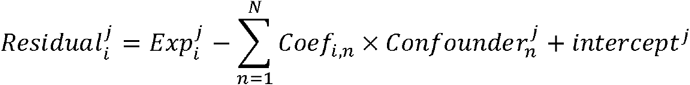

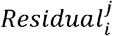 refers to the residual value of sample i gene j, 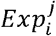 refers to the gene expression level for sample i and gene j, Coef_i,n_ refers to the coefficient of sample i for confounder n, 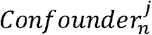 refers to the value of confounder n for gene j, Intercept^*j*^ is the intercept, the expected value of gene j when all coefficients for a sample are zero. We further applied our analysis to the residual values 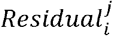.

### 2.3. Co-expression network analysis

To produce the tissue-specific modules, we applied The Weighted Gene Correlation Network Analysis (WGCNA), and the related R packages [15] were applied to calculate the co-expression networks. The WGCNA algorithm groups related genes into gene modules (networks) based on their co-expression patterns and topological similarity to neighbor genes in the modules. The algorithm calculated a similarity co-expression matrix using correlation coefficient *corci*(*i,j*) or all genes (we used the bi-weight mid-correlation measure that accounts for outliers by assigning larger weights to values closer to medians).

The co-expression matrix transforms into an adjacency matrix using the soft threshold power β, to which co-expression similarity is raised.

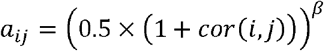

The value *a_ij_* represents the resulting adjacency that measures the connection strengths. The power *β*= 12 based on the criterion of approximating the scale-free topology of the network based on the recommendation of the authors of the WGCNA [15]package for “signed” networks, meaning that the co-expressed modules include strictly positive correlations between the nodes. We used eigengenes to calculate the inter-tissue correlation between modules, each module’s first Singular Value Decomposition (SVD) [24] component; Figure 2 demonstrates two tissues that had three modules deemed metabolic using our plurality vote mechanism. The mechanism works as follows: Approximately 300 pathways from KEGG were split into 6 categories the first six categories from of https://www.genome.jp/kegg/pathway.html. If the intersection of the genes associated with the module and the specific pathway received a Benjamin-Hochberg [25] p-value more significant than 0.05 then one vote was cast for the category associated with the pathway; otherwise nothing was added. The category that scored in absolute terms the highest was inferred to be of that category. In case metabolic score the same highest score with another category than metabolic was selected.

**Figure 2.**
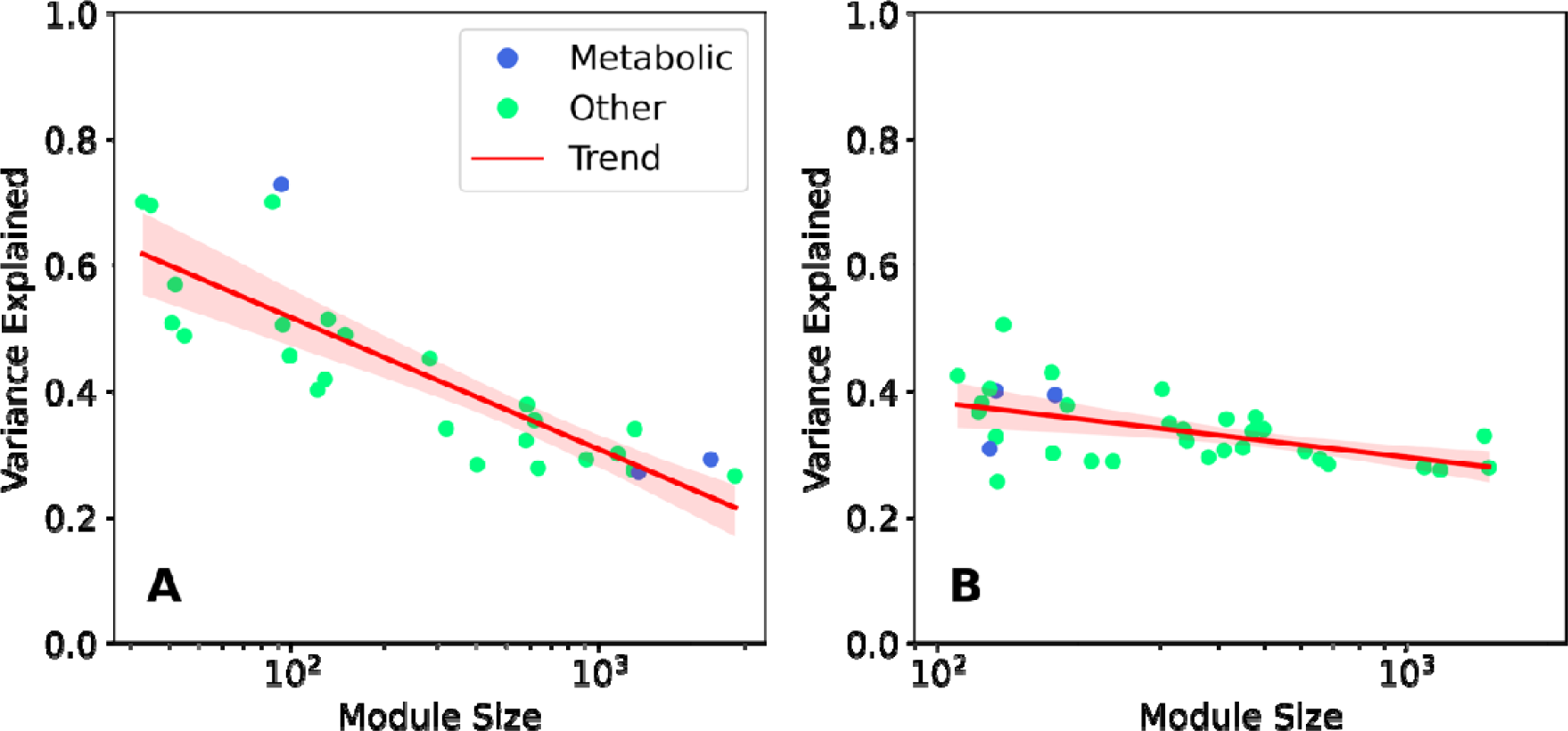
Modules annotations and variance explained by eigengenes. tissue. Each circle represents a co-expression module. The colors represent the classification of the modules specifically the blue colors are metabolic modules and the green modules were non-metabolic classifications namely: Genetic Information Processing, Environmental Information processing, Organismal, Human Disease or Unclassified. Unclassified modules that do not have a single pathway obtained an adjusted p-value of 0.05. The pathways are from the six parent categories of https://www.genome.jp/kegg/pathway.html. A. Adipose Visceral. B. Artery Aorta. Module size is number of genes in a module which is ranked in a logarithmic scale.

### 2.4. Module annotation

We inferred the module function by applying gene enrichment based on the KEGG pathways [26]and classifying each pathway into six categories. We used the KEGG PATHWAY classification [27] includes six categories. We extended into seven categories by splitting human diseases into two categories - human disease/metabolic and human disease/other. In practice, we classified the modules into one of eight categories: Metabolism, Genetic, Environmental, Cellular, Organismal, Human Diseases, Human Diseases / Metabolic, and Unclassified. We used the adjusted p-value scores (Benjamini-Hochberg [25]) of pathway enrichment for each KEGG pathway and module to classify each module according to a plurality vote we developed among the KEGG pathway categories, where each adjusted p-value < 0.05 counted as one vote.

### 2.5. Community analysis

We devised a random test to evaluate the significance of community sizes within a network of metabolic modules. The network was constructed based on the correlation between modules identified using the Weighted Gene Co-expression Network Analysis (WGCNA) algorithm. In this network, nodes represent the modules created by WGCNA, and edges represent significant correlations between these modules. Only edges with a p-value < 0.05 and correlation scores greater than 0.25 were preserved in the network. The community detection within this network was performed using the k-clique Community Partitioning Method (CPM) [16] algorithm, with a chosen value of _*k*=3._

The method began with defining a network containing only metabolic modules, with a total size of _*x*=40_. In this network, communities were identified using the CPM algorithm, and each community, denoted as *C*, was characterized by its size *S*_*c*_.

The next step involved a randomization process, where we randomly selected 40 modules from a set created in 19 different tissues, maintaining the exact proportions of only metabolic modules found in those 19 tissues. This selection included both metabolic and non-metabolic modules. This random selection was performed 1000 times, and for each randomized network, we detected communities using the same CPM algorithm. Metrics were then calculated for each community <u>*C*</u> in the randomized networks. The metrics included the Max Community Sie <u>*S*</u> , Max Community Degree <u>*d*</u>, and Average Community Degree <u>*d*</u>_*avg*_ A community *C* in the original network was judged significant if its size *S*_*c*_ was larger than the sizes of communities detected in the randomized networks.

Statistical analyses [28] were conducted using one-sample Wilcoxon signed-rank tests. First, we compared the max community sizes in the randomized networks with the specific value of <u>*S*</u> *<12* from the original metabolic network. The test provided significant statistical support for claiming that the maximum community sizes in the randomized networks are smaller than 12 (p-value: 4.34×10^-153^ ). Next, we compared the max community degrees in the randomized networks with the specific value of <u>*d*</u>*<10*, and the test showed that the max community degrees are significantly smaller (p-value: 1.62×10^-161^ ). Similarly, we compared the average community degrees in the randomized networks with the value of <u>*d*</u> <6 and found that they are significantly smaller (p-value: 1.41×10^-161^ ). We conducted the same test for networks with a correlation larger than 0.3 and k-click communities with *k*=3, and the results provided robust evidence that the max community sizes, max community degrees, and average community degrees in the randomized networks are significantly smaller than the corresponding values in the original metabolic networks. Specifically, for the new data set, the p-values were 5.02×10^-131^ , 1.79×10^-131^ and 8.98×10^-128^ for max community size, max community degree, and average community degree, respectively. The statistical significance of these findings demonstrates that the original metabolic network possesses unique structural properties not observed in the randomized networks, reinforcing the biological relevance and specificity of the detected community structures within the metabolic network.

### 2.6. Randomization tests

We used Monte Carlo randomization tests [17] to evaluate the statistical significance of our results. To construct a random distribution, we sampled modules randomly, as the number of metabolic modules, with 1000 repetitions, and counted the number of inter-modules Pearson’s [29]correlation coefficient across various cutoffs. We performed a stratified split of the random modules into tissues to confirm that we preserved the same imbalance as the metabolic modules across tissues.

2.7 Gene-level analysis

We selected 12 tissue pairs from cohort 1 by selecting three pairs from four different groups we defined – (1) tissue pairs with inter-tissue metabolic modules that ranked pairwise in the top 3 highest positive correlations (rows 1-3 Table 1), (2) tissues with inter-tissue metabolic modules that ranked pairwise in the top 3 strongest negative correlations (rows 4-6 Table 1), (3) tissue with a module of that contains a node with a high degree of edges and a tissue with weak connectivity for all metabolic modules. For example, in cohort 1, the highest degree of edge connectivity in the lung tissue is two for positive and negative correlation among the four modules. In contrast, the adipose-subcutaneous tissue has a module 038 with a degree of negative edges (rows 7-9 Table 1), (4) similar tissue types or tissues physically adjacent to each other (rows 10-12 Table 1). An example of physical adjacency is esophagus mucosa and esophagus muscularis, and an example of a similar tissue type is skin sun-exposed and skin suprapubic. We used a KEGG global metabolic pathway (01100M) containing 1250 genes as a basis for our metabolic gene set. By applying the cartesian product from each tissue-tissue pair, we obtained more than 1 million tested associations. We generated a random set of gene*i*-to-gene tuples for control from tissue *i* to tissue*j* . We used the t-test to create the significance analysis.

**Table 1.**
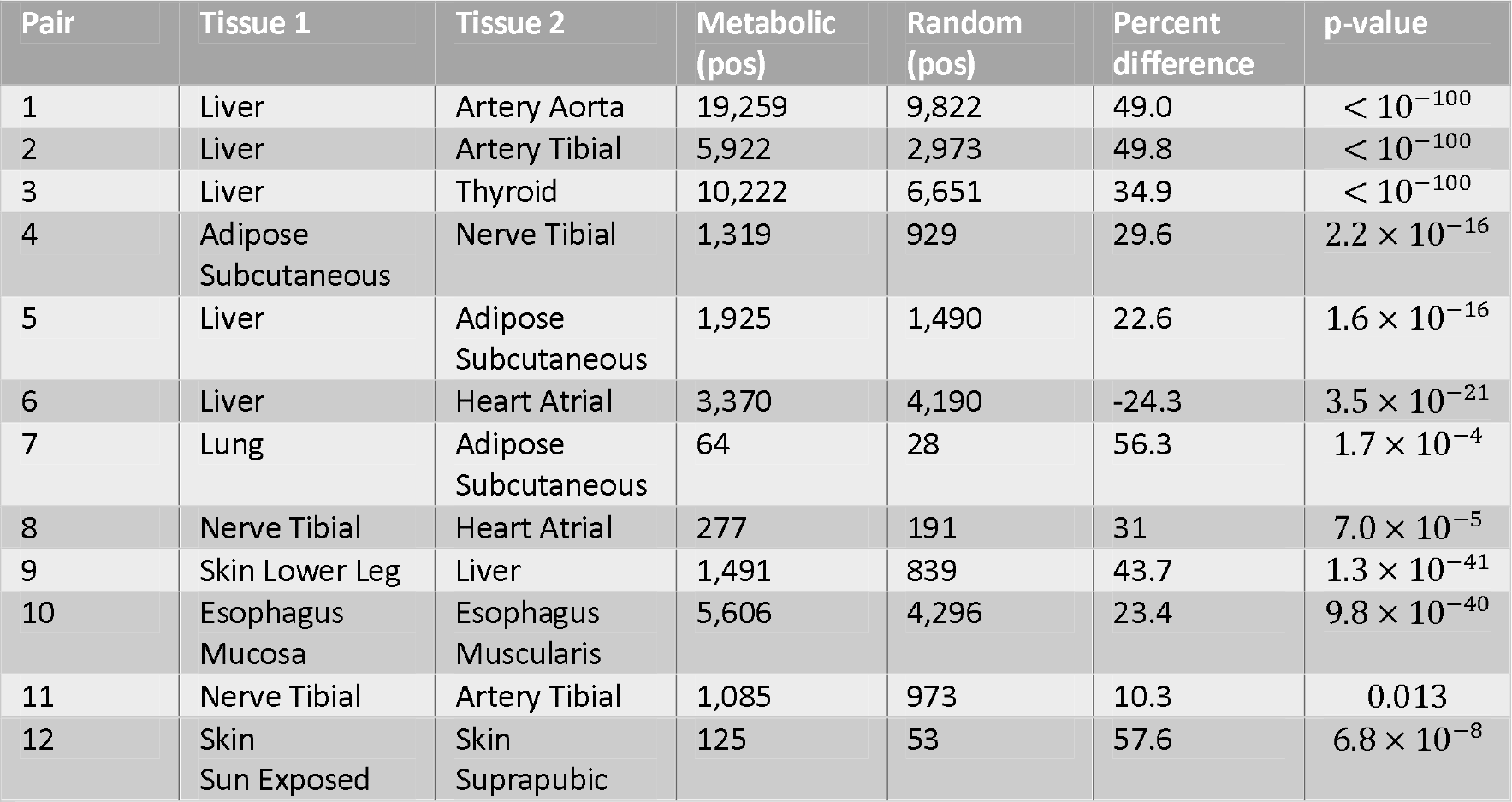
Gene-to-gene level analysis that compares the number of positive correlations (cutoff > 0.4) of more than 1 million associations between 1250 metabolic genes and randomly sampled genes generated for 12 tissue pairs.

| Pair | Tissue 1 | Tissue 2 | Metabolic (pos) | Random (pos) | Percent difference | p-value |
| --- | --- | --- | --- | --- | --- | --- |
| 1 | Liver | Artery Aorta | 19,259 | 9,822 | 49.0 | $< 10^{-100}$ |
| 2 | Liver | Artery Tibial | 5,922 | 2,973 | 49.8 | $< 10^{-100}$ |
| 3 | Liver | Thyroid | 10,222 | 6,651 | 34.9 | $< 10^{-100}$ |
| 4 | Adipose Subcutaneous | Nerve Tibial | 1,319 | 929 | 29.6 | $2.2 \times 10^{-16}$ |
| 5 | Liver | Adipose Subcutaneous | 1,925 | 1,490 | 22.6 | $1.6 \times 10^{-16}$ |
| 6 | Liver | Heart Atrial | 3,370 | 4,190 | -24.3 | $3.5 \times 10^{-21}$ |
| 7 | Lung | Adipose Subcutaneous | 64 | 28 | 56.3 | $1.7 \times 10^{-4}$ |
| 8 | Nerve Tibial | Heart Atrial | 277 | 191 | 31 | $7.0 \times 10^{-5}$ |
| 9 | Skin Lower Leg | Liver | 1,491 | 839 | 43.7 | $1.3 \times 10^{-41}$ |
| 10 | Esophagus Mucosa | Esophagus Muscularis | 5,606 | 4,296 | 23.4 | $9.8 \times 10^{-40}$ |
| 11 | Nerve Tibial | Artery Tibial | 1,085 | 973 | 10.3 | 0.013 |
| 12 | Skin Sun Exposed | Skin Suprapubic | 125 | 53 | 57.6 | $6.8 \times 10^{-8}$ |

## 3. Results

We present here the results divided into three steps of our methodology to detects a global whole-body synchronization pattern. We note that we generate step 1 and step 2 for both cohorts but chose to proceed with Cohort 1 for step 3 and further analysis since these donors were relatively healthy and not under a ventilation machine prior to death.

### 3.1. Step 1: Co-expression networks generation and annotation

We applied our methodology to 19 distinct tissues obtained from two human cohorts included in the GTEx project (see Methods). In cohort 1, we identified 40 metabolic co-expression modules out of 609, while in cohort 2, we identified 52 metabolic co-expression modules out of 652. We used the WGCNA algorithm for co-expression network analysis, Singular Value Decomposition (SVD) [22] of the generated co-expression modules for each tissue and plurality votes for metabolic modules annotation, as elaborated in the Methods section.

### 3.2. Step 2: Positive to negative metabolic transcriptomes co-regulation pattern

To evaluate the positive to negative metabolic transcriptomes co-regulation across tissues we conducted an inter-tissue module-to-modules pairwise analysis. We calculated the extent to which inter-tissue pairs of metabolic co-expression modules are positively and negatively co-regulated between the 19 tested human tissues, as opposed to randomly chosen modules. We also expected the randomly chosen actual modules to exhibit associations between various tissues. However, we wanted to evaluate the extent of a general whole-body level coordination of metabolic modules compared to the random ones. For this aim, we counted the pairwise metabolic correlation coefficients among modules, exceeding various cutoffs, and compared them to a similar random sampling of modules that are not merely metabolic. To generate a random distribution, we applied a random sample 1000 times for each cohort (see Methods). We measure our results by pairwise correlation instances, pairwise instances surpassing a p-value threshold, and by the ratios of positive to negative correlations. Figure 3 presents the results for both cohorts.

**Figure 3.**
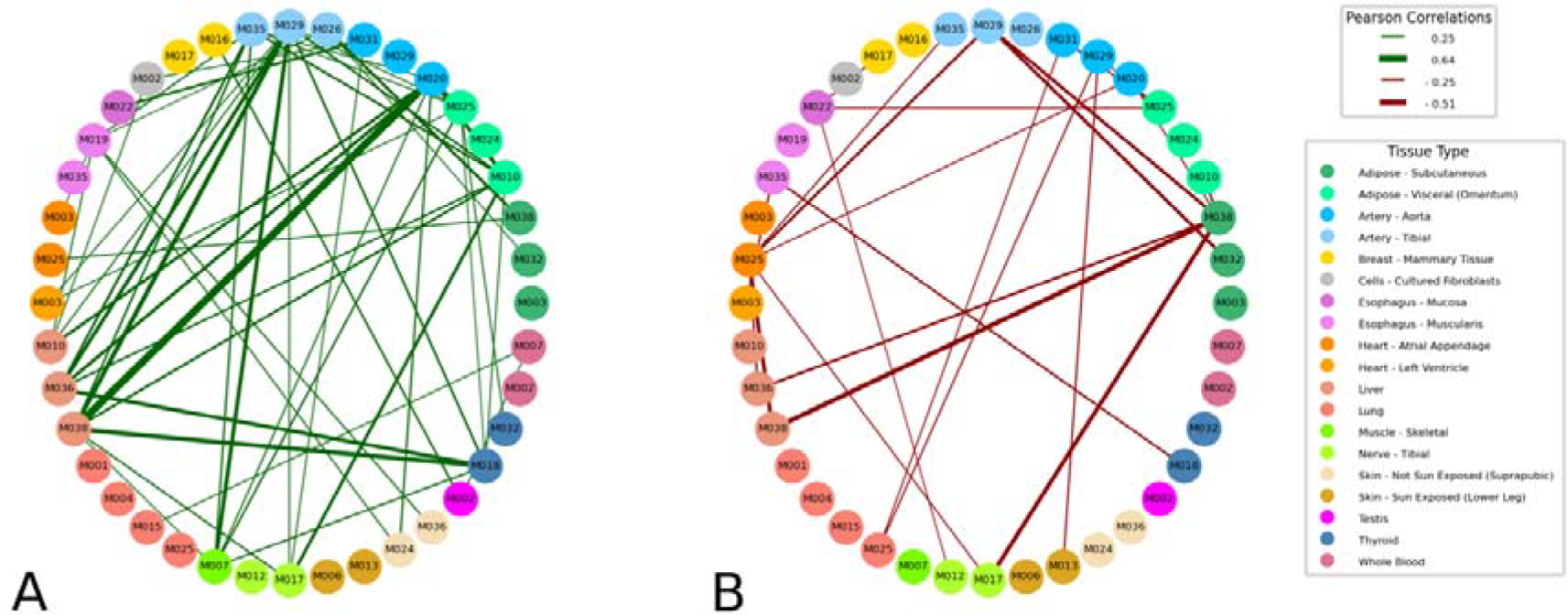
Describes inter-tissue associations found between modules. The cutoff was a Pearson correlation coefficient of ±0.25, with wider lines indicating stronger correlation coefficients. We ignored correlations within the same tissue since such correlations are generally stronger and would skew our analysis of the generally weaker inter-tissue correlations. **A**. Displays positive correlations for cohort 1. **B**. Displays negative correlations for cohort 1. Liver module 038 has the strongest positive inter-tissue correlation with Artery Aorta module 020. Liver module 038 also has the strongest negative inter-tissue correlation paired with Adipose Subcutaneous module 038. Fifty-one positive metabolic inter-tissue correlations exceed the 0.25 threshold compared to 22 negative inter-tissue correlations exceeding the -0.25 threshold. The trend that in metabolic modules, the positive correlations surpass the negative correlation per given threshold exists over a large spectrum of correlation thresholds; the 0.25 cutoff is an illustrative example. For cohort 2, the cohort with the additional influence of a mechanical ventilation device see supplemental file.

Figures 3A and 3B present the inter-tissue meta-network connectivity of the metabolic modules for positive and negative associations for Cohort 1 (edges presented for correlation coefficient cutoff of +/-0.25) between the metabolic modules across the 19 tissues in each cohort. Cohort 2 is in supplemental figure S01. Figure 4 presents the inter-tissue meta-network connectivity of the metabolic modules for positive and negative associations for Cohort 1 and Cohort 2. The number of positive inter-tissue associations (highlighted in green to the left) exceeds the negative. In addition, the liver is one of the leading tissues in node connectivity and correlation strength. In cohort 1, the metabolic liver module 038 is part of the pair with the highest positive correlation of nearly 0.64 with Artery Aorta 020 and the most significant negative correlation with Adipose Subcutaneous with a negative correlation level over -0.5 in magnitude.

**Figure 4.**
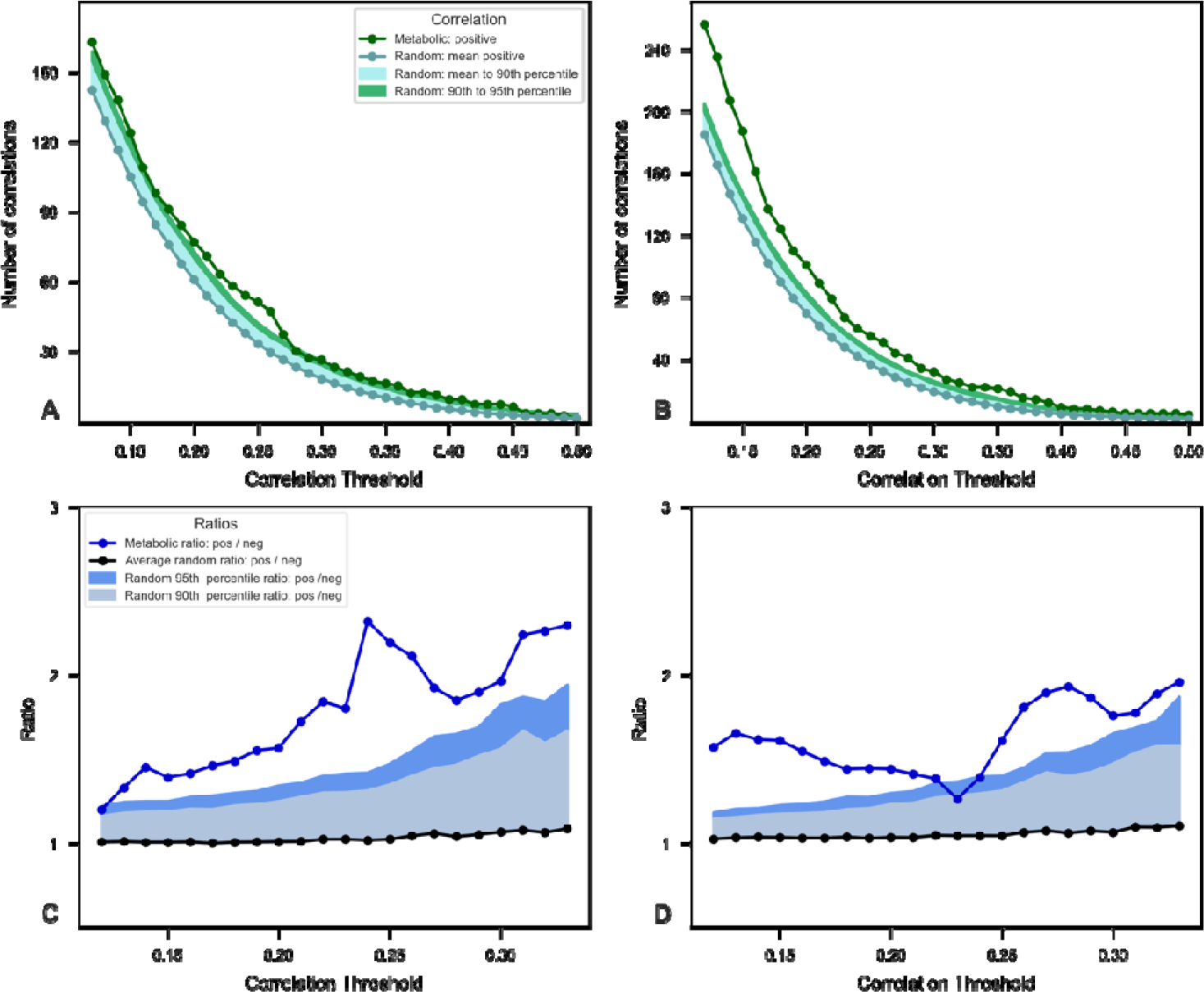
Pairwise inter-tissue association counts. We applied this analysis to two cohorts. Cohort 1 had 609 modules, of which 40 were deemed metabolic modules and a potential of 751 pairwise inter-tissue association counts. Cohort 2 had 652 modules, of which 52 were deemed metabolic and a potential of 1257 inter-tissue modules. The comparison was between correlations found per threshold of the inter-tissue metabolic modules to a random set of selected modules. The random selection process was performed 1000 times to form a distribution, and we compared the metabolic modules. A. Cohort 1 comparison of metabolic pairs exceeding a given correlation threshold to the random group. B. Cohort 2 comparison of the metabolic modules to its associated random group. C. Cohort 1 ratio of positive to negative correlation exceeding a threshold compared to the ratio of positive to negative counts in the random group. D. Cohort 2 ratio of positive to negative correlation exceeding a threshold compared to the ratio of positive to negative counts in the random group.

### 3.2. Step 3: Cross-tissue synchronization of metabolic transcriptomes

In this step we conducted a meta-network-level analysis to detect a global cross-tissue positive synchronization pattern of metabolic transcriptomes. For this we used community analysis utilizing the CPM (Clique Percolation Method) algorithm [16]. A community is a group of nodes (here a node is an eigengene and an edge represents connectivity measure by Spearman correlation) that is densely interconnected. We evaluated the system-level whole-body metabolic positive co-regulation testing the statistical significance of metabolic community size and connectivity measures as opposed to randomly sampled modules (Fig. 5). For this, we generated the community of metabolic modules across the 19 tissues and 1000 communities of randomly chosen modules (see methods). The analysis in Figure 5 shows that the metabolic community includes 12 modules (nodes) and spans across seven different tissues, Adipose Visceral, Artery Aorta, Artery Tibial, Liver, Muscle Skeletal, Nerve Tibial, and Thyroid (Fig. 5A), and that metabolic transcriptomes form a significantly larger community (p-value < 0.05) with higher maximal and average modules connectivity when compared to control.

**Figure 5.**
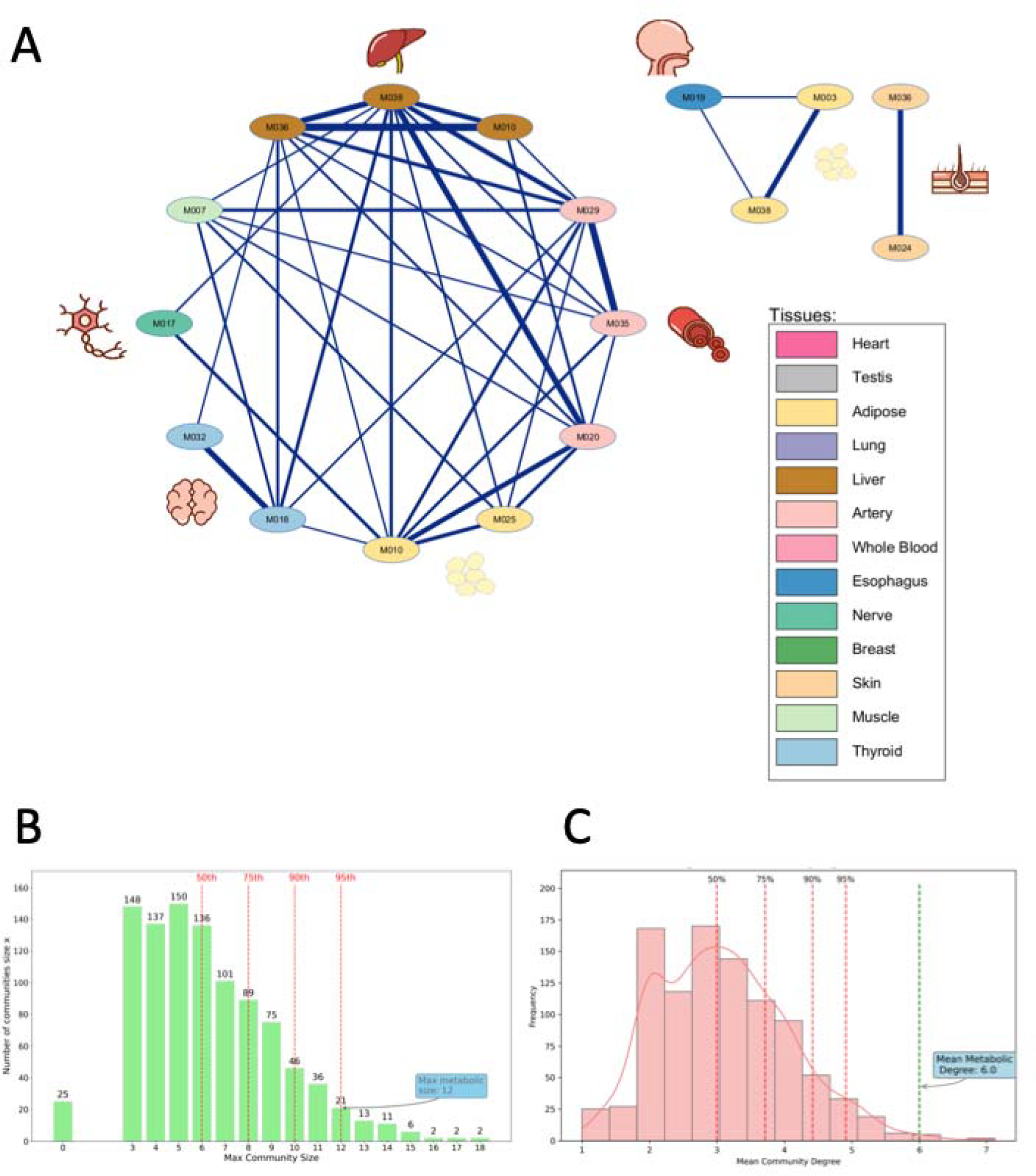
Community analysis of the network of metabolic co-expression modules across 19 tissues for cohort 1 A. Community of metabolic modules, k = 3. The nodes represent metabolic modules in tissues. The edges represent Spearman’s correlation coefficient between the nodes (edge=1 if correlation coefficient > 0.25, adjusted p-value < 0.05) B. Distribution of mean community degree for the random sample and the measure for the metabolic community. C. Distribution of maximal community nodes degree for a random sample and the measure for the metabolic community. D. Distribution of maximal community size for the random sample and the measure for the metabolic community. The inter-tissue metabolic modules form a significantly larger community size and are densely connected, exceeding the 95th percentiles for the various measures.

## 4. Gene-level coordination analysis

For validation purposes, we conducted an inter-tissue gene-to-gene level analysis. For this we selected 12 representative tissue pairs (see Methods) and used 1250 genes as a basis for our metabolic gene set derived from KEGG global metabolic pathway (01100M). By applying the cartesian product from each tissue-tissue pair, we obtained more than 1 million gene*i*-to-gene*j* tuples for each test and control. We used the t-test to compute the significance analysis. Table 1 summarizes the results, presenting the number of gene-gene pairs at correlation thresholds of 0.4 Pearson’s correlation coefficient for all 12 tissue-tissue pairs, the statistical significance of the difference between the gene-level positive correlations counts, and the random sample counts. Additional supplementary plots for all 12 tissue pairs show similar trends for correlation thresholds of 0.3, 0.4, and 0.5 and are presented in supplemental figure S5. Table 1 demonstrates significantly smore robust inter-tissue connectivity in 11 of 12 pairwise comparisons of metabolic genes as opposed to randomly chosen (11 out of 12 binomial p-value = 5.3 x 10-S). Moreover, the one exception was from the group that displayed the inter-tissue module pairs with the most negative correlations.

## 5. Discussion

In this research, we developed a new bioinformatic framework to observe a systems-level synchronization pattern of metabolic transcriptomes across human tissues. Our framework analyzes the meta-network of metabolic co-expression modules across 19 tissues in two human cohorts. We generated 609 modules for cohort 1 and 652 modules for cohort 2 across the tissues and annotated them. We then used dimensionality reduction representation of the modules to test the metabolic positive labeled assortativity [19] of our whole-body network, i.e., to test if similar modules (metabolic modules) exhibit more robust positive connectivity and larger communities. Labeled assortativity is the tendency in networks that nodes with the same label will have greater interconnectivity. We finally performed a gene-level validation for 12 pairs of tissues.

Our approach demonstrates a whole-body positive co-regulation of metabolic transcriptomes in tissues and a significantly high ratio of positive-to-negative associations. Furthermore, we show that tissues’ metabolic transcriptomes are densely and positively connected and form larger communities than randomly chosen modules (Fig. 4). Our analysis demonstrated that when observing the associations cumulatively, we detect a clear pattern of whole-body positive coordination and coregulation of metabolic transcriptomes. When we compared metabolic modules to a comparable pool of random modules, we noticed that the number of positive associations between cross-tissue metabolic modules significantly exceeded (p-value < 0.05 for most cutoffs, Fig. 3A, and 3B) the random modules in Cohort 2 and approached the level of statistical significance in Cohort 1. The smaller sample size of Cohort 1 explains the difference in statistical significance. Furthermore, the positive to negative metabolic module ratio is statistically significant for both cohorts (p-value < 0.05 for most cutoffs, Fig. 3C and 3D) compared to 1000 collections of comparable random modules. Furthermore, for Cohort 1, we also checked at the gene-to-gene level. We obtained that metabolic genes exhibit greater coordination patterns than a random set of genes statistically significant at a statistically significant level beyond a p-value of 0.01.

When analyzing the networks, we detected that the liver tissue is a component in inter-tissue module pairs with the highest correlation rates. Prior studies have confirmed this observation [10], [30], [31]. The liver was an outlier in our “positive” pattern for Cohort 2 and exhibited multiple negative associations with distant tissues. It is worth noting that in Cohort 2, representing the donors that were ventilated prior to their death, Esophagus Muscularis and Lung modules exhibited a high degree of inter-tissue synchronization. Esophagus Muscularis module number 053, jointly with Nerve Tibial 033, share the most significant positive correlation. The two Esophagus Muscularis share many connections with other tissues. The prominence of Esophagus Muscularis may be due to a technical factor: the mechanical ventilation machine, which is physically nearby.

Our approach detected positively labeled assortativity of inter-tissue metabolic modules in both cohorts. This technique can serve as a tool for detecting coordination patterns for multiple organisms and pattern changes across disease conditions. However, because inter-tissue correlations inherently have low correlations, and a dimension reduction approach compounds this, we recommend using this technique for comparison purposes when over 100 pairwise samples are available for each condition to make further comparisons. Our observation on sample sizes explains why, in Fig 3B, the results exceeded were more significant at the lower correlation threshold than in Figure 3A. See supplemental Figure S4 for inter-tissue counts per cohort.

## Supporting information

see supplemental file.

## Author contributions

J.S. conceived the research idea and methodology. E.Z.S. implemented the code and refined the methods in practice. D.M. performed the community analysis. E.Z.S. and J.S. analyzed the results. J.S , E.Z.S. and D.M. wrote the paper.

## Supplementary data

Supplementary data is available in the supplementary file.

Conflicts of interest: None declared.

## Data availability

The GTEx data used is freely accessible at https://gtexportal.org/home/, computer code for project is found at https://github.com/pelikanGIS/BioninformaticsJune2023

