## Supplementary material for "Whole-body positive synchronization pattern of metabolic transcriptomes": see supplemental file.

### Figure S1

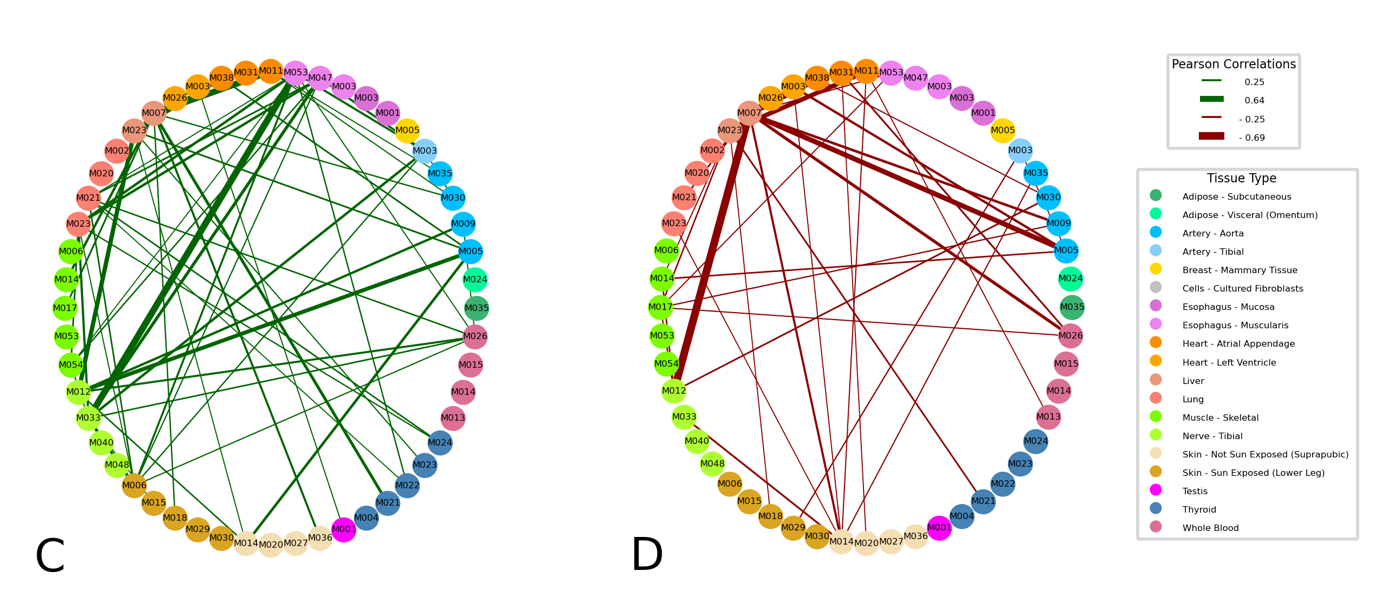

**Figure S2:** Describes inter-tissue associations found between modules. The cutoff was a Pearson correlation coefficient of ±0.25, with wider lines indicating stronger correlation coefficients **C.** Displays positive correlations for cohort 1. **D.** Displays negative correlations for cohort 1. As with Figure 3A and 3B from Cohort1 the positive associations exceed the negative ones. In addition to the Liver tissues composing a disproportionately large proportion of inter-tissue correlations we observed that the Lung, Esophagus Muscularis, and Nerve Tibial also add many edges to the graph.

### Figure S2

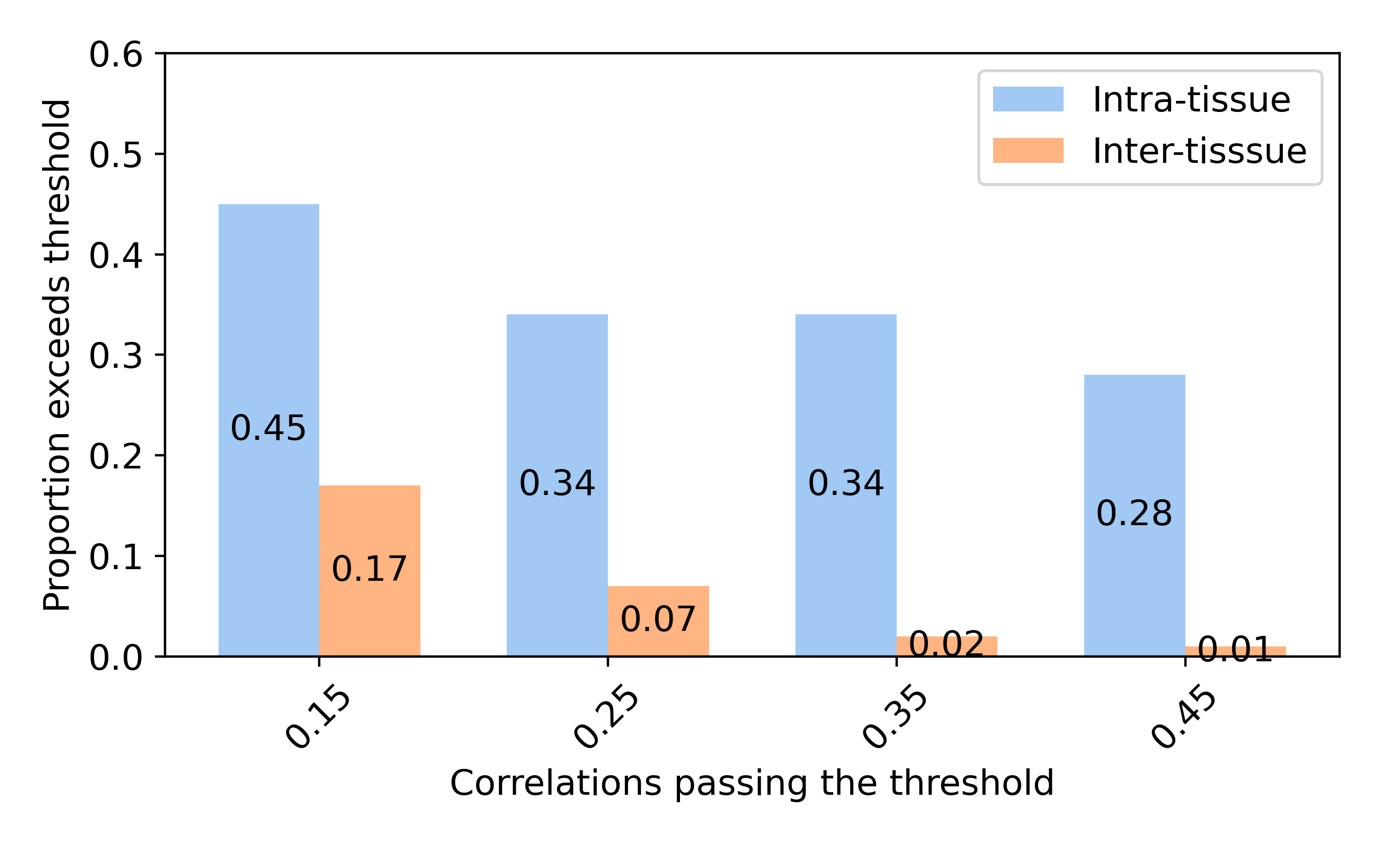

**Figure S2:** Proportion of pairwise modules exceeding a threshold by network type.

In total we had 40 modules in cohort 1 that were deemed as Metabolic. The potential pool of pair-wise correlations can be calculated using the hand shake formula. $\frac{n(n-1)}{2}$. In this instance $n=40.$ Thus we have a pool of 780 potential correlations that can exceed a given threshold. However, we are interested in inter-tissue correlation and a tissue can have multiple metabolic modules, the purely inter-tissue association modules comprised 751 of the 780 metabolic pair-wise association, leaving 29 modules as intra-tissue.

The graph above shows that approximately 45% of the 29 (see Figure 2 Supplemental) metabolic module pairs that were intra-tissue or exceed a correlation threshold of 0.15.

### Figure S3

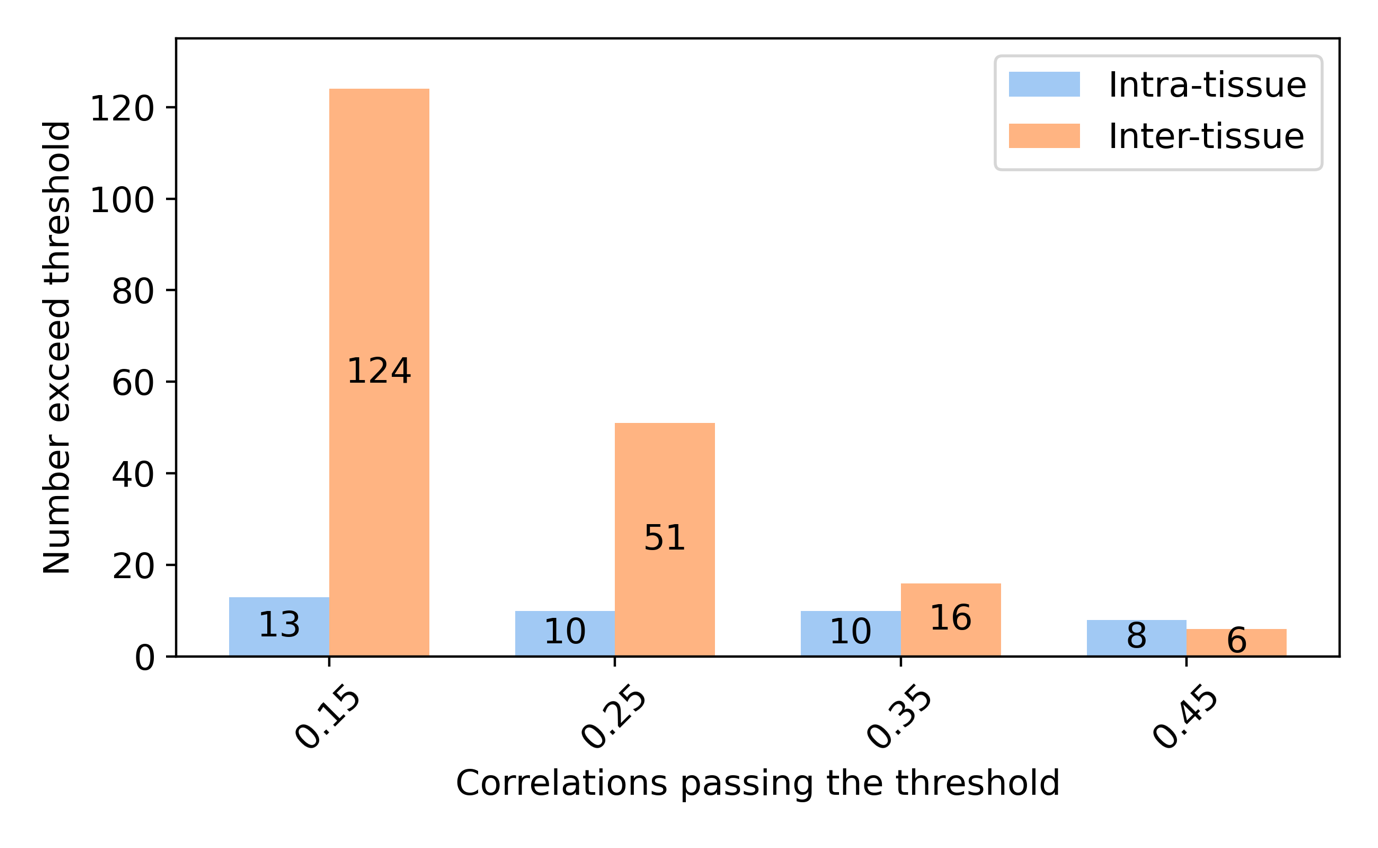

**Figure S3:** Number of pairwise modules exceeding a threshold by network type.

The actual number was 13 of 29 (See Figure 3 Supplemental) . We observe that for lower correlation threshold the including intra-tissue modules may not impact the analysis much, however when we analyze higher correlation we achieve a situation that the although the 29 modules of the 780 are less than 4% of all pairwise correlations, the make the absolute majority of instances passing a given threshold.

### Figure S4

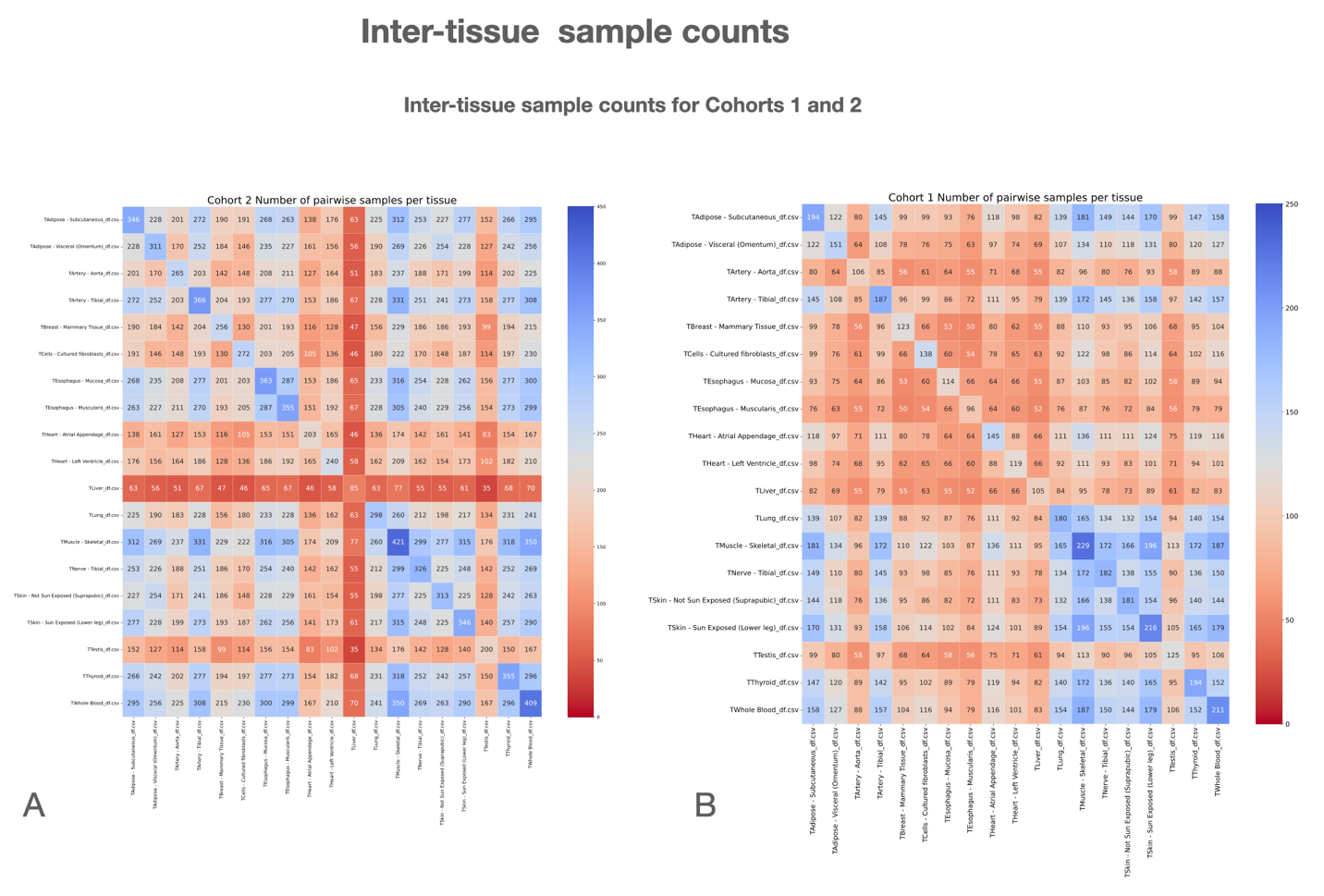

**Figure S4:** Display the actual samples from the same donor of different tissues. Along the main diagonal we see the number of samples in a specific tissue. **A.** Shows the number of pairwise samples in cohort 2 and **B.** shows the number of samples in cohort 1. For example, in cohort 1 we has 194 samples of Adipose Subcutaneous tissue, and 229 sample from the Muscle – Skeletal tissue. If we look at the row of Adipose – Subcutaneous and the column of Muscle – Skeletal, we observe that in the intersection of both groups we had 181 sample we could include for pairwise comparisons between the two groups.

### Figure S5

**Figure S5:**

Gene to gene ratios for metabolic. The figures that start with 1 namely 1A, 1B, 1C are from the group that included an inter-tissue metabolic module with the strongest positive correlations. The figures that start with 2 namely 2A, 2B, 2C are from the group that included an inter-tissue metabolic module with the strongest negative correlations. The figure that start with 3 namely 3A,3B,3C are from the group of tissues had included a tissue with a metabolic module that had a high degree of connectivity and a tissue that all the all metabolic modules had low degrees of connectivity at correlations threshold of 0.25. The figures that start with 4 namely 4A, 4B, 4C are tissues that have a similar function or a near one another in the human body.

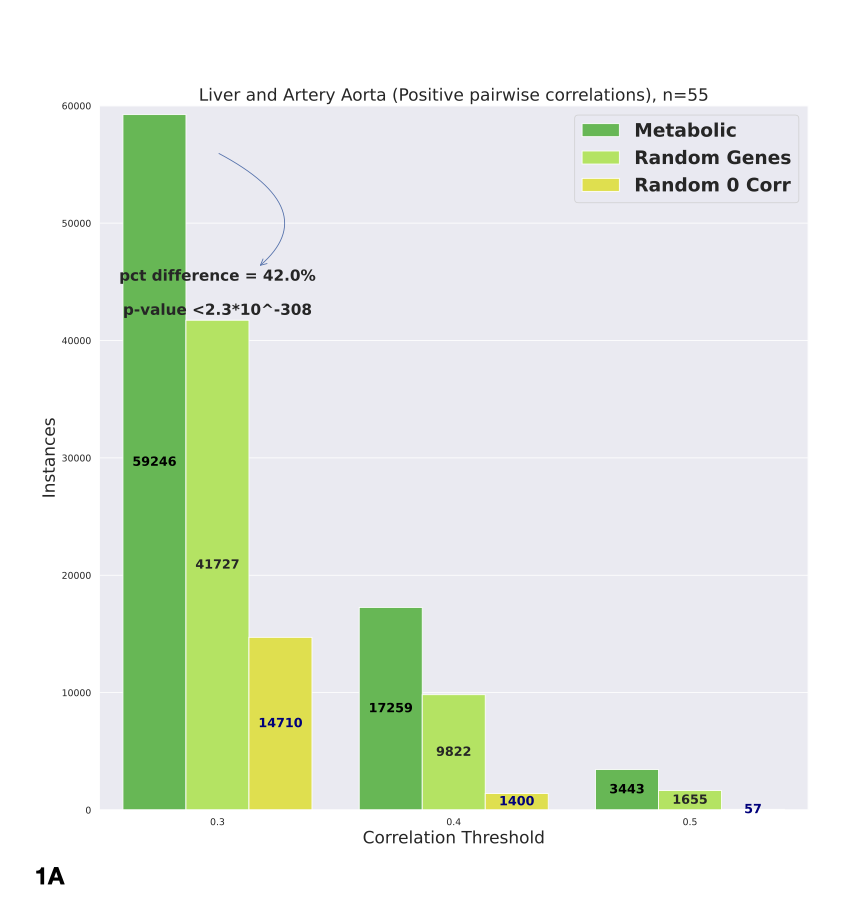

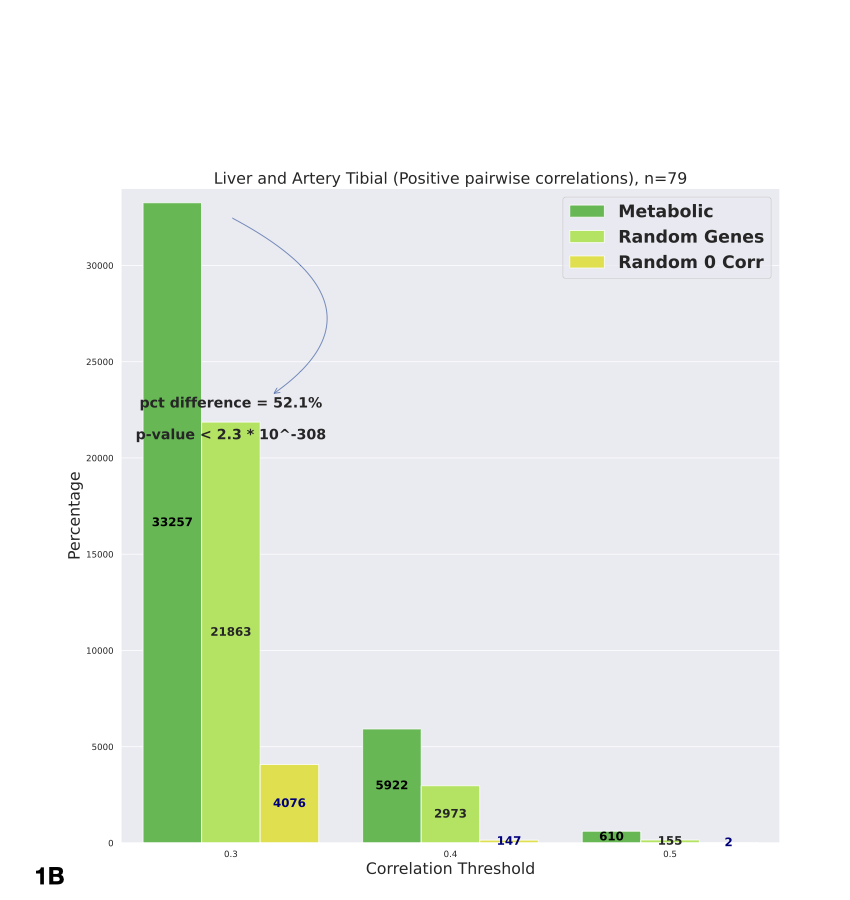

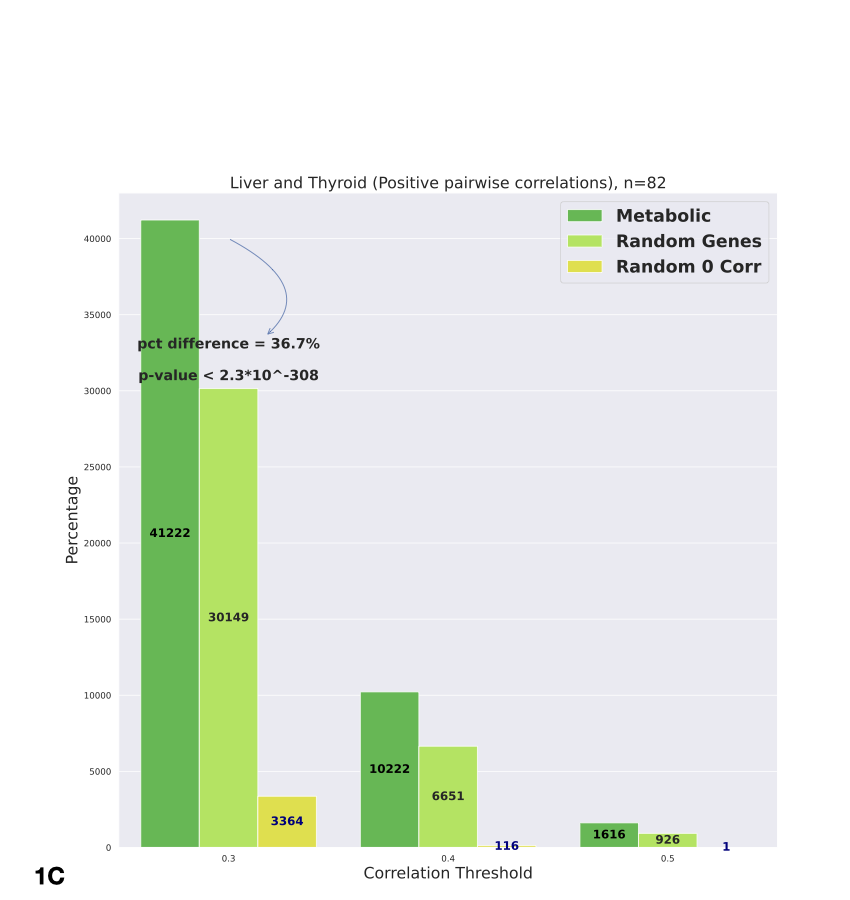

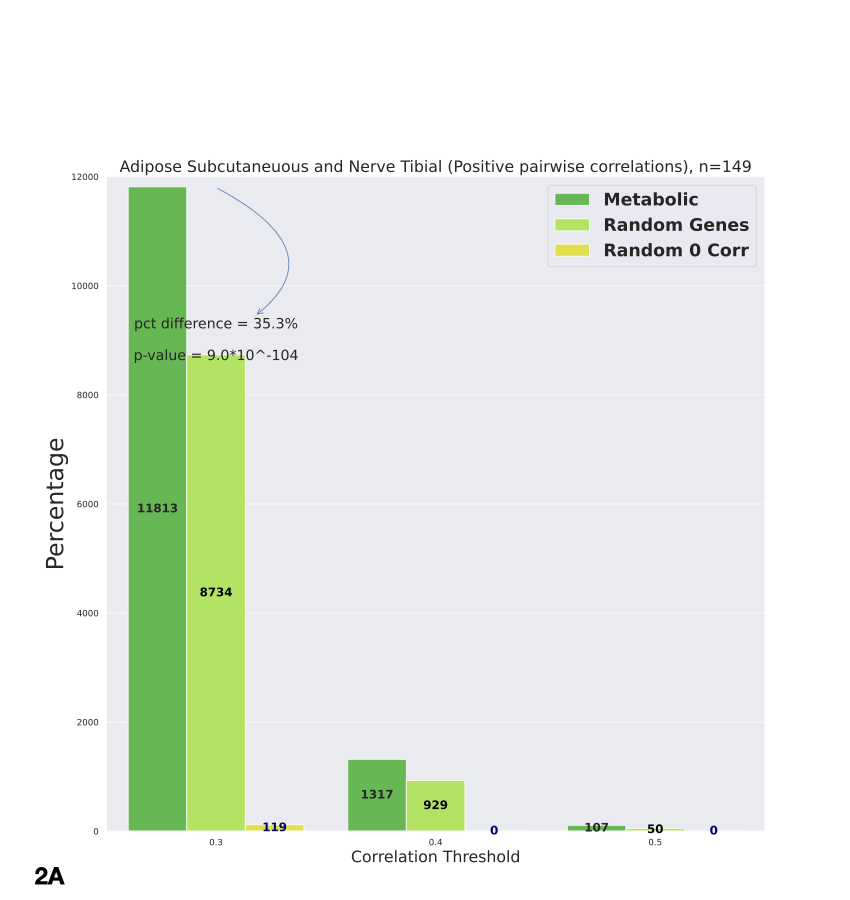

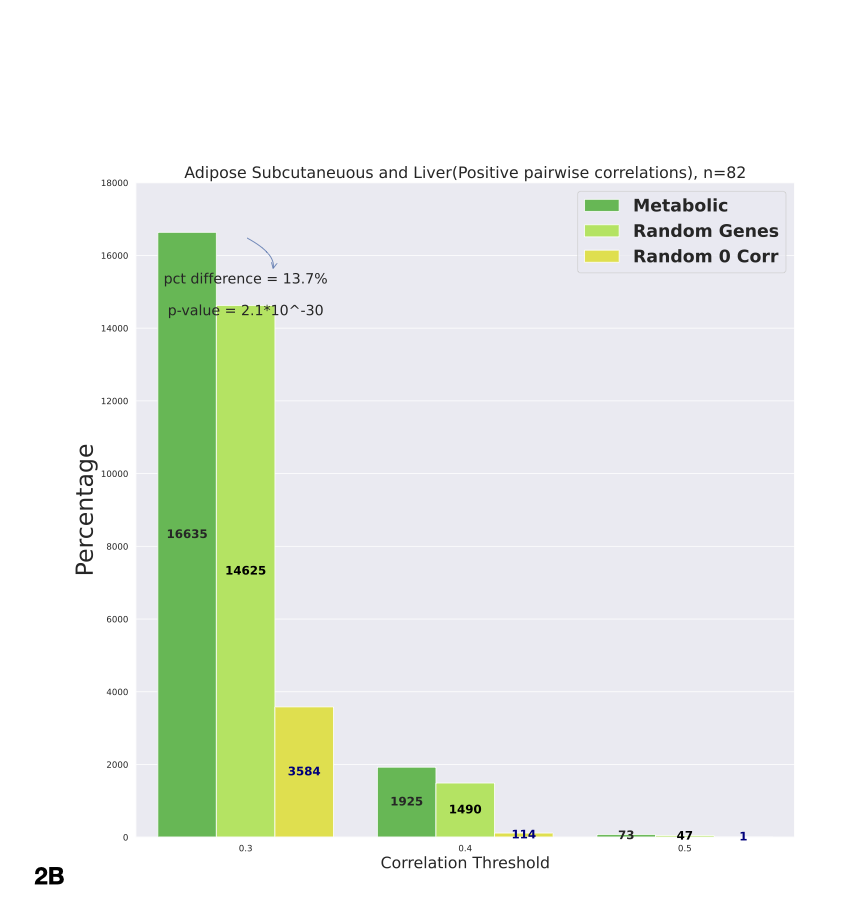

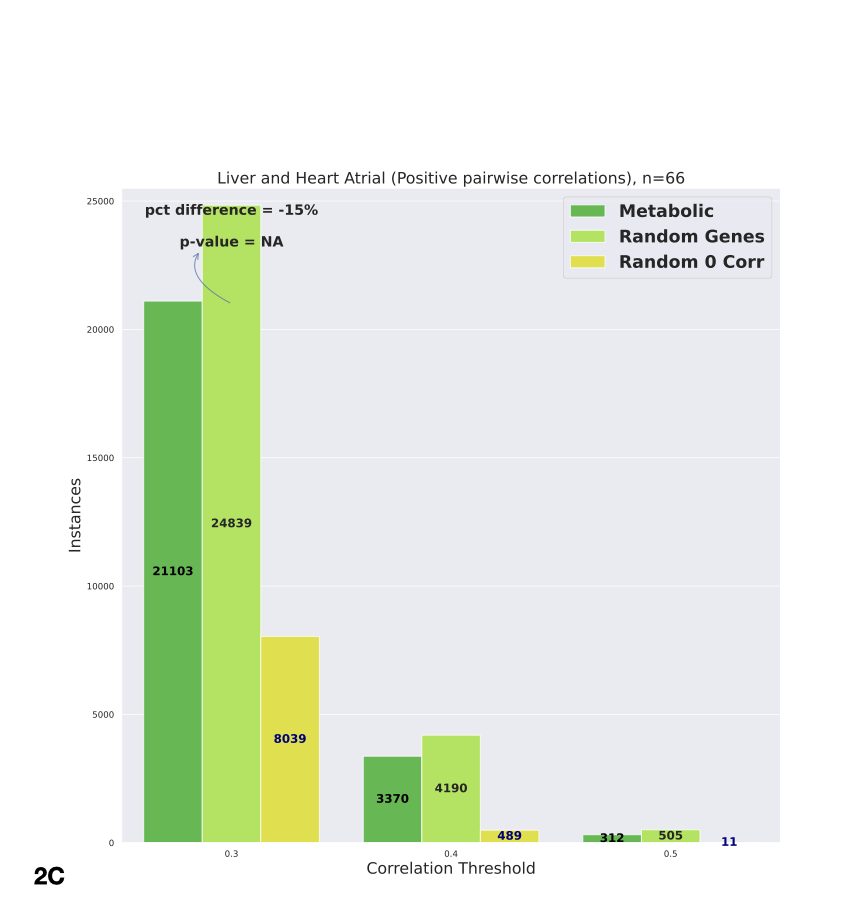

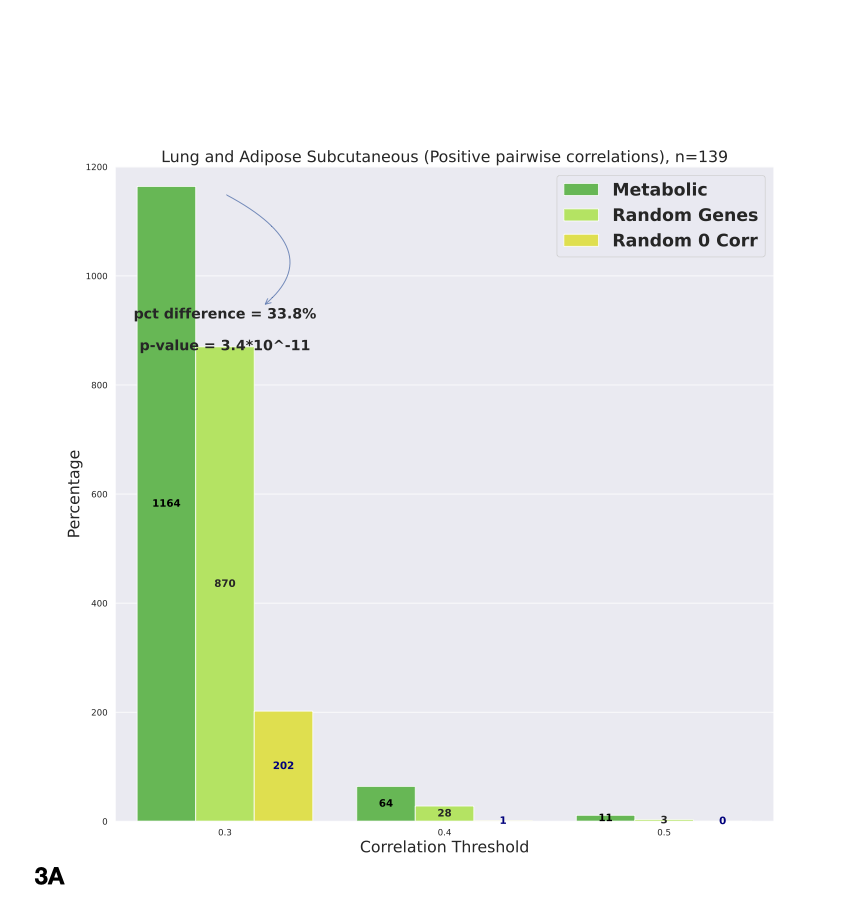

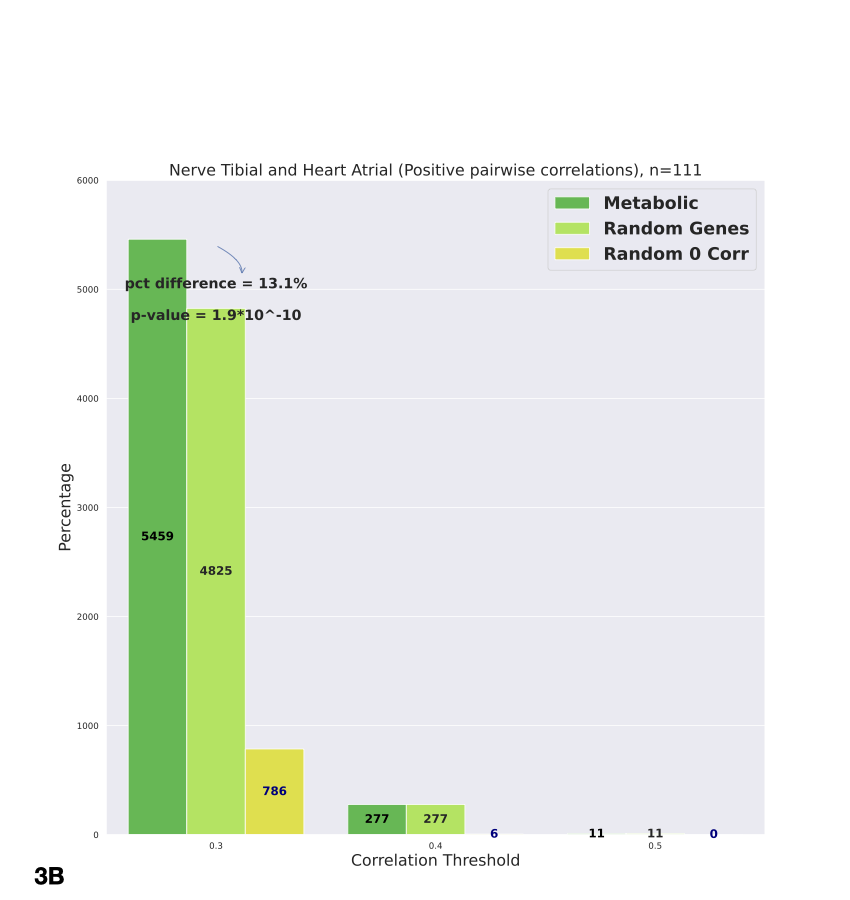

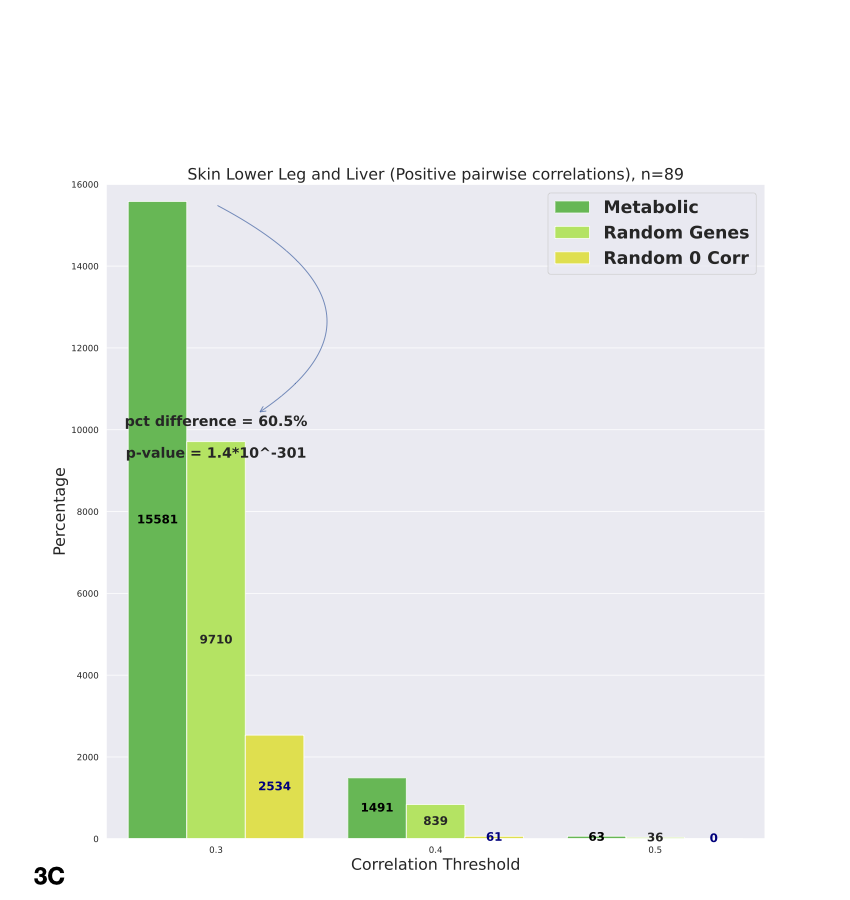

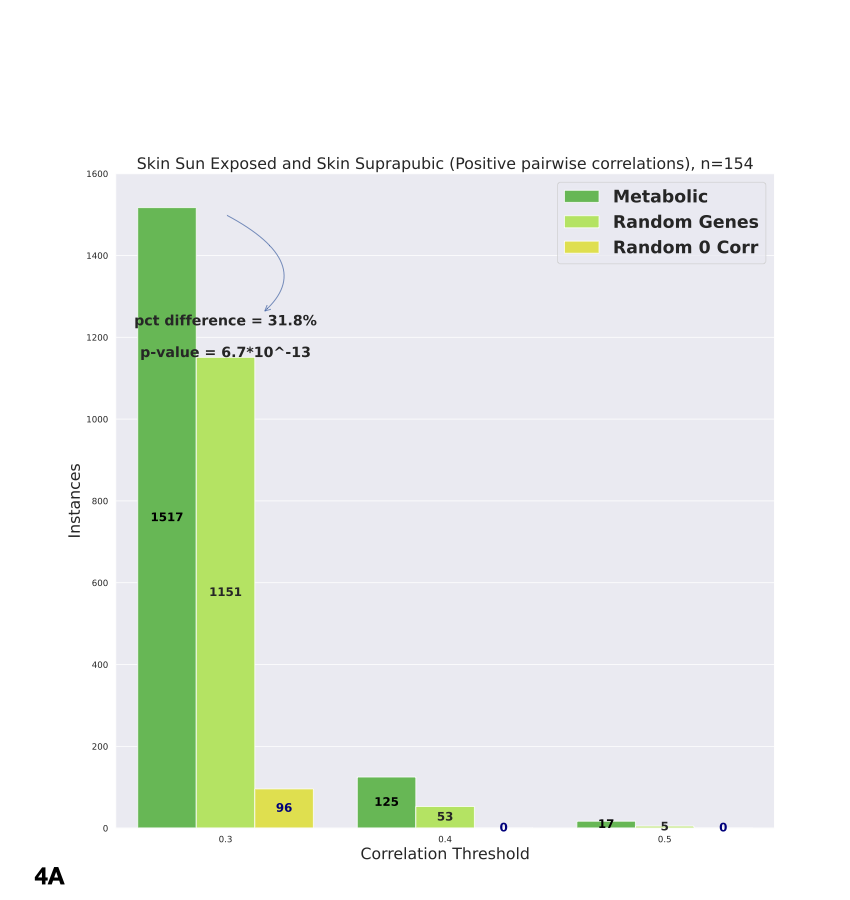

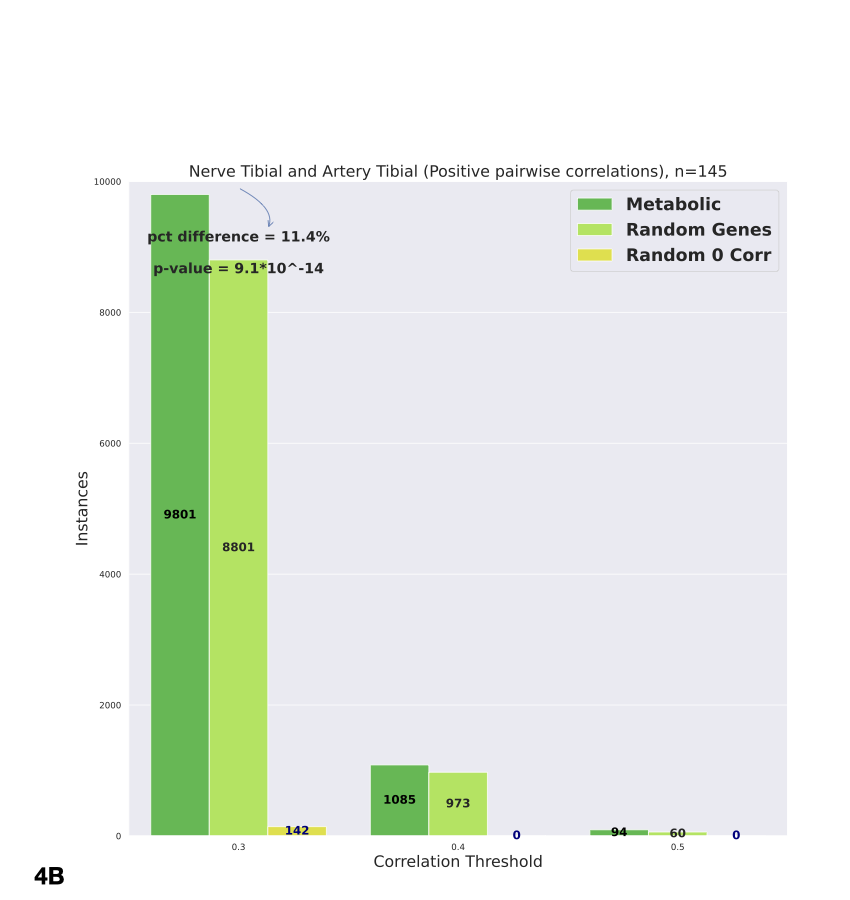
